# Tensorflow Based Deep Learning Model and Snakemake Workflow for Peptide-Protein Binding Predictions

**DOI:** 10.1101/410928

**Authors:** Gokmen Altay

## Abstract

In this study, we first present a Tensorflow based Deep Learning (DL) model that provides high performances in predicting the binding of peptides to major histocompatibility complex (MHC) class I protein. Second, we provide the necessary Python codes to run the model and also easily input large train and test peptide binding benchmark dataset. Third, we provide Snakemake based workflow that allows to run all the model and performance analysis over all the different test alleles at once in parallel over computer and clusters. We also provide comparison analysis of the performances of various models. Finally, in order to help attaining to the best possible DL model by a community effort, this work is intended to be a ready to modify base model and workflow for the global Deep Learning community with no domain knowledge in MHC-peptide binding problem and thus provides all the necessary reference code templates and benchmarking data sets for further developments on the presented model architecture. All the reproducible Python codes, Snakemake workflow and benchmark data sets and a tutorial are available online at https://github.com/altayg/Deep-Learning-MHCI.

## Introduction

Cytotoxic T-cell lymphocytes (CTLs) interacts with complexes of peptides and major MHC class I (MHCI) molecules presented on the cell surface to detect and destroy the cells that harbor intracellular threats [1]. Accurate prediction of bindings between peptides and MHC protein helps the design of more potent, peptide-based vaccines and immunotherapies [2].

There are many machine learning (ML) methods that were developed to infer the binding between peptides and MHC class I protein with high accuracy [2-7]. Among them, NetMHC [1] and NetMHCpan [3], which are neural network (NN) based models, are considered as the state of the art models to predict binding between peptides and MHC class I molecules [7]. Deep Learning [8] is an advanced version of NN, where the word deep mainly refers to the deeper number of hidden layers in the model. DL based methods were shown to be outperforming classical ML methods where there is very large data available and thus found applications in various areas such as image recognition [9], speech recognition [10], predicting the activity of potential drug molecules [11], analyzing particle accelerator data [12], reconstructing brain circuits [13], and predicting the effects of mutations in non-coding DNA on gene expression and disease [14] and many others [8]. There have been very recently several DL based approaches developed to predict binding between peptides and MHC class I molecules, such as MHCflurry [7], MHCnuggets [4], HLA-CNN [2] and DeepMHC [6]. In this study, we present a convolutional neural network (CNN) based DL approach and demonstrate that, on average, it outperforms state-of-the art algorithm NetMHCpan [3] and also provides competitive performances with some of the other contemporary algorithms that have open source software code available to perform comparisons. In order to ease the readability of the manuscript, we name our DL model as DL-MHC in the rest of the paper.

### Motivation

Although there are many available models to predict MHCI binding prediction, there is still no consensus on the best performing and optimal model on the problem. Although, most of the new models are published by presenting slightly better performances than netMHCpan [3], it is still considered as the state-of-the-art [7]. Most of the Deep Learning (DL) based models have similar performances to each other. Although, each publication with a new DL model shows a slightly better performance than some others, there is no reproducible model that was shown to provide better performances in general with comparison to all the other available DL models considering most of all the alleles. Each related publication comes with slightly different set of alleles and thus, performances are valid for the studied alleles of the presented models. Also, the tiny differences in the average performances would not mean that the presented model would perform best in general. Especially, when it comes to Deep Learning, randomization has a significant role in its performances. At each run, the models might perform slightly different on the same dataset. An attempt to set all the randomizations as constant in the model might also limit the performance of the models on average as there are many different alleles to consider. Among others, these might be some of the reasons that there is still no consensus on a best performing DL model for MHCI and peptide binding prediction problem and NetMHC [1] and netMHCpan [3] are still considered as gold standards and referenced usually as the state-of-the-art.

We consider that the best performing DL model architecture might be attained by a global community effort. For example, after an overwhelming interest by the global Machine Learning (ML) community on the famous MNIST dataset (yann.lecun.com/exdb/mnist), it is very recently reported that a novel model was developed that provides 0% error rate [15]. However, applications of protein-peptide binding predictions with DL or ML is being developed only by a number of researchers with the domain knowledge. This study also aims engaging the global DL community in this very important problem of immune system. By this challenge, we expect that very skilled DL developers, with no domain knowledge, will eventually come up with dramatically best performing model for the peptide-protein binding prediction problem. In order to encourage DL community in general, we specified some potential obstacles and provided solutions.

One reason that might prevent the wider DL community not to engage with this important problem is that there is no easily run workflow available, such as Snakemake [16], to run a newly developed DL model over all the different alleles at once. Entering amino acid sequences of peptide data into DL model is another related challenge. In this study, we provide large peptide train and test datasets with the necessary Python source code to easily access and use them in any new application easily along with the documentation to explain them to any user with no background about the problem. We also provide a step-by-step tutorial to easily run all the presented DL models and the Snakemake workflow to evaluate the performance of all the different alleles at one run in parallel on a single machine making use of multiple CPUs or submit as batch jobs over clusters.

Another reason that might prevent the wider DL community not to engage with the problem might be that there are not much open source DL models that use the most popular DL library Tensorflow (tensorflow.org). Developed by Google, Tensorflow dominates the field with the largest active community, which has around three times as many GitHub forks and more than six times as many Stack Overflow questions than the second most popular framework [17]. It also returns by far the highest number of resulting pages compared to any other DL library when searched by any internet search engine. Tensorflow allows developing any kind of DL architecture and not limited with its ready to use modules. Most of the experienced DL developers use or know Tensorflow and thus an open source exemplary model along with already implemented very large protein-peptide binding dataset would help them easily engage with the problem. This way, even though they do not have any domain knowledge, they can start modifying the provided running DL model architecture and utilizing their long experience in DL model development, they might end up with dramatically higher performance scores than the reference exemplary model. In this study, we developed a Tensorflow based DL model, named as DL-MHC, which on average provides competitive performances with the state-of-the-art model netMHCpan [3]. However, as finding the best model is not the main purpose of this study, we do not claim that it performs best considering all the DL models available. As we present in the performance results, DL-MHC provides better results than netMHCpan [3] in some of the alleles and similar performances to other DL models and there are not very large significant differences among them in general.

A similar study approach to accelerate the development of DL methods within biology by providing application examples and ready to apply and adapt code templates was recently published in [18]. It provides exemplary DL models and source codes for the prediction of subcellular localization, protein secondary structure and the binding of peptides to MHC Class II molecules. However, they used somewhat unpopular NN Lasagne library based on Theano DL library (www.deeplearning.net/software/theano). In 2017, it was announced that Theano developer team will put an end to Theano development after the 1.0 release and thus is regarded currently as being almost dead. On the other hand, in this study, we used the most popular DL library, Tensorflow, and showed that the provided reference DL model, DL-MHC, is providing competitive prediction performance with the state-of-the-art model NetMHCpan in the prediction of binding of peptides to MHC Class I molecules. Although there are several DL implementations on the problem, to the best of our knowledge, only one of them provide a Tensorflow based open source DL model [19], which uses Recurrent Neural Networks (RNN) and can be utilized as complimentary with our CNN based DL model by the DL community. Also, it does not provide any comparison with other similar models. In this study, but also provide a comparison analysis with various model which can be used as a quick performance reference table by the developers who might want to compare the performance of their new models quickly by the provided model performances on the given large benchmark dataset. This study uniquely provides Snakemake based workflow that allow running all the models and performance analysis over all the different test alleles at once in parallel over computer or clusters. This would save a significant amount of time for developers and let them focus only on the model developments.

As the summary of main contributions of this study, first, we provide a Tensorflow based DL model that provides high prediction performances and can be used as a base to easily modify and attain to best performing DL model architectures. We also provide very large peptide-protein binding train and test datasets along with their Python source codes ready for using to input them in any new DL model. We provide comparison analysis over various recent models with the used benchmark datasets that can be used as a quick reference performance scores table for any newly developed DL model. Second, we provide a Snakemake based workflow that allow running all the models and performance analysis over all the different test allele datasets at once in parallel over computer or clusters. We also provide a step-by-step tutorial for running the presented DL model and the Snakemake workflow. We hope these contributions will help more engagement of the global DL community with no domain knowledge on the binding of peptides to MHC proteins and thus provide an infrastructure to further developments. This way, we might help the problem being better aware of wider developers and cause getting wider attention to it.

## Methods and Materials

### Dataset

The first dataset we used is the benchmark dataset used in the analysis of HLA-CNN [2] and we downloaded the dataset from the GitHub repository link given in the paper for the software of HLA-CNN. This dataset was filtered, processed and prepared by [5], which includes HLA class I binding data curated from four popular publicly available MHC datasets that are SYFPEITHI [20], AntiJen [21], MHCBN [22] and IEDB [23]. As described in [2], the peptides that contain unknown or indiscernible amino acids, denoted as ‘X’ or ‘B’, are removed from the dataset before the training. The test datasets for 15 different alleles were downloaded from IEDB automatic server benchmark page (http://tools.iedb.org/auto_bench/mhci/weekly/). In the dataset, binding is classified if ic50 measurements are less than 500nm. This training and test dataset is called as Dataset 1 in this study in the repository to make the modifications easier in future if new and different datasets are wanted using with the workflow. We provide this dataset in our GitHub repository of DL-MHC but it is already available in HLA-CNN repository [2]. Using the exact same dataset with HLA-CNN study allowed us to directly compare DL-MHC with the comparison analysis of HLA-CNN [2]. This way, we make sure that we compare with the exact same results produced by the authors of the algorithms that we compare and avoid any potential usage bias on this dataset.

### Encoding Peptide Sequence for Deep Learning Model Input

The best way to explain the procedure is by example. In Table 1, we illustrate how we encode a peptide sequence ‘KAYKSIVKY’ into a matrix of 9 rows and 20 columns. The exemplary peptide sequence has 9 amino acids and thus 9-mer in length.

**Table 1:** Example of encoding the peptide sequence ‘KAYKSIVKY’ into one-hot encoded data matrix.

|  |  |  |  |  |  |  |  |  |  |  |  |  |  |  |  |  |  |  |  |
|---|---|---|---|---|---|---|---|---|---|---|---|---|---|---|---|---|---|---|---|
| K | 0 | 0 | 0 | 0 | 0 | 0 | 0 | 0 | 1 | 0 | 0 | 0 | 0 | 0 | 0 | 0 | 0 | 0 | 0 |
| A | 1 | 0 | 0 | 0 | 0 | 0 | 0 | 0 | 0 | 0 | 0 | 0 | 0 | 0 | 0 | 0 | 0 | 0 | 0 |
| Y | 0 | 0 | 0 | 0 | 0 | 0 | 0 | 0 | 0 | 0 | 0 | 0 | 0 | 0 | 0 | 0 | 0 | 0 | 1 |
| K | 0 | 0 | 0 | 0 | 0 | 0 | 0 | 0 | 1 | 0 | 0 | 0 | 0 | 0 | 0 | 0 | 0 | 0 | 0 |
| S | 0 | 0 | 0 | 0 | 0 | 0 | 0 | 0 | 0 | 0 | 0 | 0 | 0 | 0 | 1 | 0 | 0 | 0 | 0 |
| I | 0 | 0 | 0 | 0 | 0 | 0 | 1 | 0 | 0 | 0 | 0 | 0 | 0 | 0 | 0 | 0 | 0 | 0 | 0 |
| V | 0 | 0 | 0 | 0 | 0 | 0 | 0 | 0 | 0 | 0 | 0 | 0 | 0 | 0 | 0 | 0 | 0 | 1 | 0 |
| K | 0 | 0 | 0 | 0 | 0 | 0 | 0 | 0 | 1 | 0 | 0 | 0 | 0 | 0 | 0 | 0 | 0 | 0 | 0 |
| Y | 0 | 0 | 0 | 0 | 0 | 0 | 0 | 0 | 0 | 0 | 0 | 0 | 0 | 0 | 0 | 0 | 0 | 0 | 1 |

The procedure to encode a peptide sequence into a one hot encoded data matrix is as follows. We first assign a unique integer number to each one of the 20 possible amino acids.

All possible amino acids are ACEDGFIHKMLNQPSRTWVY. From left to right we assign their indexes as unique numeric representations instead of each amino acid letter. In the same order:

[0, 1, 2, 3, 4, 5, 6, 7, 8, 9, 10, 11, 12, 13, 14, 15, 16, 17, 18, 19]. Namely, the index of A is 0, the index of C is 1 and the index of Y is 19. In order to represent A as one-hot encoded vector, we generate a one-hot encoded vector of size 20 with all zero values except the first index, as it is set to be 1 like this [1 0 0 0 0 0 0 0 0 0 0 0 0 0 0 0 0 0 0 0]. As seen in the above example from Table 1, for a 9-mer length peptide sequence, we repeat the process for each amino acid and build a matrix of 9 rows and 20 columns. If the peptide sequence was 11, then we would generate a data matrix of 11 rows and 20 columns in a similar way. Similar approach for encoding peptide sequence is also used in [19]. In our DL model, the Python function that performs this mapping operation is named as *Pept_OneHotMap.*

### Proposed Deep Learning Model, DL-MHC

We build a Deep Learning model with Convolutional Neural Network [8]. As seen in Figure 1, Our module architecture has three parallel connections of different filter sizes, which was inspired by the Inception module from GoogleNet [24]. We call our **D**eep **L**earning (DL) model for **MHCI** binding prediction as DL-MHC for the rest of the paper. It has 8 of two-dimensional (2D) convolutional neural network (CNN) layers in each of the three parallel connections, which are then concatenated to flatten before entering into the fully connected dense layer of 100 nodes. The output is passed through a dropout process with 50% keep rate and enters to final layer with 2 nodes for the two possible output labels, binding and not-binding.

**Figure 1.**
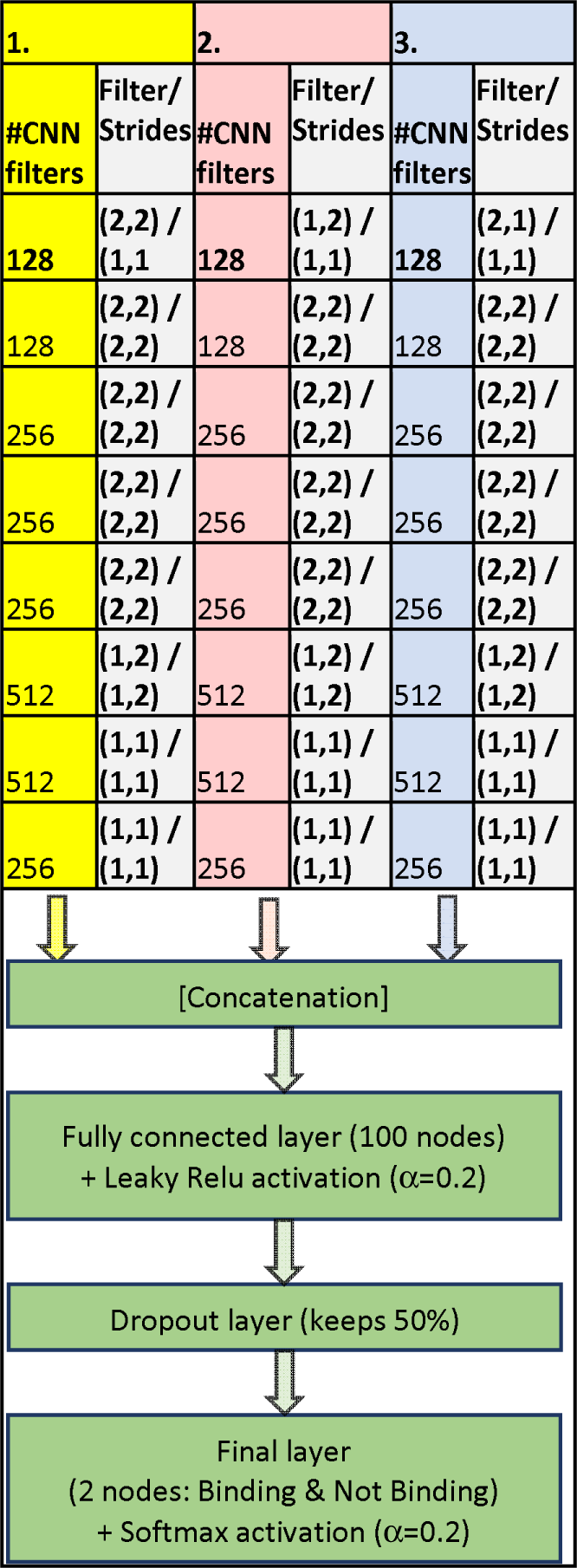
Deep Convolutional Neural Network Architecture: The data enters from connections 1, 2 and 3. CNN filters are two-dimensional (2D) convolutional filters. There are three parallel connections, 8 layers each, of CNN filters. In each of the 2D convolutional layer of each of the three parallel connections, Leaky Relu activation (α=0.2) is used.

We use AUC (Area Under the ROC (receiver operating characteristic) Curve) as the main performance metric because it is mostly the convention to use AUC in the related publications. As we selected AUC, following the tradition to measure the performances, we have used Softmax activation function in the output layer. If we had selected Accuracy or Precision as the performance metric, we would use Sigmoid function as the final activation function. For the optimization of the cost function, Adam optimizer [25] is used. We also employed adaptive learning rates as input to the optimizer. The learning rate starts with 0.003 and decreases with iterations until 0.0001 at the least minimum. The maximum number of iterations (epocs) are set as 2000 for the current analysis of the manuscript. The batch size for the mini batches is set to 40. We did not use the max-pooling as it did not help to the performances in our analysis. However, with a different DL architecture, this process might be helpful and suggested for trying in the future developments.

### Early Stopping Criteria

We developed a slightly different approach than others as early stopping criteria to the problem of interest. To our knowledge, other peptide-protein binding prediction NN based models use a single validation dataset. If the performance score (e.g. accuracy) or validation loss does not improve after a predefined number of epocs, then the model stops training even though it does not loop through the maximum number of epocs defined before training. In DL-MHC, instead of a single validation dataset, we use three different small validation datasets (20 samples each) and define a stopping rule based on the improvements of their three accuracy scores together. Details of it can be seen in DLMHC.py script available in the repository. If there is no improvement in the combined performance of the validation datasets after 300 epocs, then the training is terminated. Validation datasets are randomly extracted from the training dataset, and this way, we might have more stable results by not relying on a single validation dataset for early stopping. Regarding our many trials, it is worth mentioning that this approach was observed having a significant role in the higher performance of DL-MHC. Since our study is not aiming to find the best DL model but aims paving a smooth way in that endeavor, we did not fine tune each component of our DL model. For instance, as an exemplary suggestion for a potential further improvement on this early stopping criteria might be selecting balanced validation datasets with respect to the ratio of binding or non-binding labels instead of randomly selecting them. This and similar improvements are available to try by the interested DL developers upon our proposed model and benchmarking.

## Results

We followed the general practices in most of the publications in this field and thus used AUC (Area under the ROC Curve.) and SRCC (Spearman’s Rank Correlation Coefficient) as performance metrics. We present the performance results in the below tables.

The average performances of some of the popular algorithms in Dataset 1 were presented in Table 2. Since Dataset 1 is the same dataset used in [2], we present the same performance scores for NetMHCpan, sNebula [5], and HLA-CNN from [2]. We compare those results with our proposed DL-MHC model. Regarding AUC, DL-MHC provides the best average performance compare to the others. We do not include the scores of HLA-CNN to the performance comparison because it was mentioned in [6] that HLA-CNN software code is using the test dataset as if it is a validation set during training, which is not a correct approach regarding real ML practice. This might help HLA-CNN model to have an incorrect advantage while deciding when to stop training and might lead to overestimated prediction performance. In fact, in [6] they attempted to correct this point in HLA-CNN model code and in their analysis, the resulting AUC performance of the corrected HLA-CNN model software was 2.4% lower than the previous one. Therefore, the presented HLA-CNN’s average AUC score 0.855 is approximated to be 0.834 in Table 2. This makes our proposed DL-MHC model having the highest AUC average score with 0.853. We could not include the DL model, DeepMHC, of [6] as they did not provide their open source code in the their publication. Regarding SRCC, as seen in Table 2, DL-MHC performs highest among all the others in Dataset 1 on average.

**Table 2:** Average performances of various algorithms over 14 test data sets of 9 different alleles of Dataset 1. (HLA-CNN model was shown to have lower performances than the scores quoted in this table; because its current software uses training data for stopping criteria, which is against ML practice.)

| Dataset 1 |  |  |
| --- | --- | --- |
|  | AUC | SRCC |
| DL-MHC | 0.853 | 0.559 |
| NetMHCpan | 0.794 | 0.542 |
| sNebula | 0.749 | 0.462 |
| HLA-CNN | 0.855 | 0.551 |

More specifically, as seen in Table 3, DL-MHC provides the best AUC performance on 8 out of the 14 test data sets of Dataset 1. As explained above, HLA-CNN is just for reference but is not directly compared here. The second-best performer is NetMHCpan with 5 out of 14 test datasets. As presented in Table 4, regarding the detailed SRCC scores, DL-MHC provides the highest performances in 7 out of 14 test data sets in Dataset 1. Then, the second-best performer is NetMHCpan with 5 out of 14 test data sets. Individual test set results assure the high performance scores of DL-MHC observed on average.

**Table 3:** Average AUC performances of various algorithms over 14 test data sets of 9 different alleles of Dataset 1. (HLA-CNN model was shown to have lower performances than the scores quoted in this table; because its current software uses training data for stopping criteria, which is against ML practice.)

| **Dataset 1** |  | Peptide length | NetMHCpan | sNebula | HLA-CNN | **DL-MHC** |
| --- | --- | --- | --- | --- | --- | --- |
| IEDB | HLA |  | AUC | AUC | AUC | AUC |
| 1029125 | B*27:05 | 9 | 0.959 | 0.959 | 0.918 | 0.929 |
| 1029061 | B*57:01 | 9 | 0.943 | 0.575 | 0.807 | 0.727 |
| 1028928 | A*02:01 | 9 | 0.955 | 0.909 | 0.955 | 0.909 |
| 1028928 | B*07:02 | 9 | 1 | 0.9 | 1 | 1 |
| 315174 | B*27:03 | 9 | 0.893 | 0.607 | 1 | 1 |
| 1028790 | A*02:01 | 9 | 0.574 | 0.778 | 0.681 | 0.691 |
| 1028790 | A*02:01 | 10 | 0.677 | 0.704 | 0.589 | 0.742 |
| 1028790 | B*02:02 | 9 | 0.713 | 0.68 | 0.804 | 0.708 |
| 1028790 | A*02:03 | 9 | 0.696 | 0.629 | 0.746 | 0.755 |
| 1028790 | A*02:03 | 10 | 0.75 | 0.697 | 0.837 | 0.908 |
| 1028790 | A*02:06 | 9 | 0.77 | 0.848 | 0.819 | 0.806 |
| 1028790 | A*02:06 | 10 | 0.768 | 0.68 | 0.92 | 0.864 |
| 1028790 | A*68:02 | 9 | 0.806 | 0.713 | 0.909 | 0.920 |
| 1028790 | A*68:02 | 10 | 0.62 | 0.813 | 0.991 | 0.991 |
|  | **Average** |  | **0.794** | **0.749** | **0.855** | **0.853** |

**Table 4:** Average SRCC performances of various algorithms over 14 test data sets of 9 different alleles of Dataset 1.

| **Dataset 1** |  | Peptide length | NetMHCpan | sNebula | HLA-CNN | DL-MHC |
| --- | --- | --- | --- | --- | --- | --- |
| IEDB | HLA |  | SRCC | SRCC | SRCC | SRCC |
| 1029125 | B*27:05 | 9 | 0.751 | 0.752 | 0.684 | 0.700 |
| 1029061 | B*57:01 | 9 | 0.612 | 0.169 | 0.443 | 0.284 |
| 1028928 | A*02:01 | 9 | 0.57 | 0.539 | 0.57 | 0.693 |
| 1028928 | B*07:02 | 9 | 0.648 | 0.522 | 0.648 | 0.852 |
| 315174 | B*27:03 | 9 | 0.657 | 0.179 | 0.837 | 0.836 |
| 1028790 | A*02:01 | 9 | 0.615 | 0.505 | 0.58 | 0.280 |
| 1028790 | A*02:01 | 10 | 0.407 | 0.432 | 0.327 | 0.334 |
| 1028790 | B*02:02 | 9 | 0.582 | 0.372 | 0.426 | 0.288 |
| 1028790 | A*02:03 | 9 | 0.539 | 0.477 | 0.373 | 0.522 |
| 1028790 | A*02:03 | 10 | 0.208 | 0.419 | 0.307 | 0.566 |
| 1028790 | A*02:06 | 9 | 0.63 | 0.51 | 0.578 | 0.503 |
| 1028790 | A*02:06 | 10 | 0.572 | 0.525 | 0.638 | 0.602 |
| 1028790 | A*68:02 | 9 | 0.534 | 0.482 | 0.581 | 0.660 |
| 1028790 | A*68:02 | 10 | 0.272 | 0.591 | 0.722 | 0.714 |
|  | **Average** |  | **0.542** | **0.462** | **0.551** | **0.559** |

## Conclusion

We presented a high-performance DL model, DL-MHC, comparisons with some other models, Python source code of the model and data input process of the benchmark datasets, a Snakemake based workflow to run performance analysis over all the different alleles at once in parallel. This study is aimed to provide a base DL model and benchmark datasets with all the necessary codes upon which new and better DL models can be conveniently tried and developed by the DL community who might not necessarily have the domain knowledge. Therefore, with this study, we aim to make it convenient to any DL developer to focus on this very important problem of immunology. We hope that this study will increase the amount of interest and consequently result better DL models on the problem, which might help attaining a new state-of-the-art model that performs dramatically better than current models that are approximately providing similar performances on average.

## Acknowledgement

We thank to the authors of [2] for their prompt supports to run their software. We also thank to Sandeep K. Dhanda for the discussions on MHC-peptide binding problem and its applications.

